# VPAC: Variational projection for accurate clustering of single-cell transcriptomic data

**DOI:** 10.1101/523993

**Authors:** Shengquan Chen, Kui Hua, Hongfei Cui, Rui Jiang

## Abstract

**Background:** Single-cell RNA-sequencing (scRNA-seq) technologies have advanced rapidly in recent years and enabled the quantitative characterization at a microscopic resolution. With the exponential growth of the number of cells profiled in individual scRNA-seq experiments, the demand for identifying putative cell types from the data has become a great challenge that appeals for novel computational methods. Although a variety of algorithms have recently been proposed for single-cell clustering, such limitations as low accuracy, inferior robustness, and inadequate stability greatly impede the scope of applications of these methods.

**Results:** We propose a novel model-based algorithm, named VPAC, for accurate clustering of single-cell transcriptomic data through variational projection, which assumes that single-cell samples follow a Gaussian mixture distribution in a latent space. Through comprehensive validation experiments, we demonstrate that VPAC can not only be applied to datasets of discrete counts and normalized continuous data, but also scale up well to various data dimensionality, different dataset size and different data sparsity. We further illustrate the ability of VPAC to detect genes with strong unique signatures of a specific cell type, which may shed light on the studies in system biology. We have released a user-friendly python package of VPAC in Github (https://github.com/ShengquanChen/VPAC). Users can directly import our VPAC class and conduct clustering without tedious installation of dependency packages.

**Conclusions:** VPAC enables highly accurate clustering of single-cell transcriptomic data via a statistical model. We expect to see wide applications of our method to not only transcriptome studies for fully understanding the cell identity and functionality, but also the clustering of more general data.

## Background

Single-cell RNA-sequencing (scRNA-seq) has emerged as a revolutionary tool to reveal previously unknown heterogeneity and functional diversity at a microscopic resolution. Since the first protocols were published in 2009 [1], a massive expansion in method development has derived scRNA-seq technologies with distinct advantages and applicability [2]. For example, Smart-seq2 [3] and MARS-seq [4] is preferable when quantifying transcriptomes of fewer cells, while Drop-seq [5] is preferable when quantifying transcriptomes of large numbers of cells with low sequencing depth [6]. Advances in scRNA-seq technology have resulted in a wealth of studies aiming to reveal new cell types [7, 8], assess tissue composition [4, 9, 10], identify gene regulatory mechanisms [11, 12], investigate cell development or lineage processes [13-15], and many others. With the exponential growth of the number of cells profiled in individual scRNA-seq experiments, there is a demand for novel analysis methods for this new type of transcriptomic data, which has not only much greater scale of datasets than that of bulk experiments but also various challenges unique to the single-cell context [16].

A key advantage of scRNA-seq is that it can be used to identify putative cell types using unsupervised clustering, which is essential to fully understand the cell identity and functionality. A variety of algorithms have recently been proposed for single-cell clustering. For example, CellTree produces tree structures outlining the hierarchical relationship between single-cell samples using a novel statistical approach based on document analysis techniques [17]. Other statistical approaches based on Dirichlet mixture model are shown to be well suited for single cell clustering, especially for data as unique molecular identifiers (UMI) matrix. For example, DIMM-SC models UMI count data and characterizes variations across different cell clusters via a Dirichlet mixture prior [18]. Para-DPMM further improves the clustering quality by introducing a Dirichlet process prior to automatically infers the number of clusters from the dataset, and a split-merge mechanism is adopted to improve convergence and optimality of the result [19]. Given the high dimensionality of single-cell data, clustering directly on the original dimension may affect the performance due to the intrinsic noise of single-cell data, and usually demands tedious preprocessing such feature selection, which may restrict the robustness of clustering. A simple idea is to use traditional methods to reduce dimensions and then cluster. However, there are too many alternative combinations that make it difficult for us to make a choice. Methods combining dimension reduction with classic clustering such as t-Distributed Stochastic Neighbor Embedding (t-SNE) with K-means [8], and principal component analysis (PCA) with hierarchical clustering [20] for single-cell clustering are proposed. Combining PCA with K-means, a consensus clustering approach is proposed to achieve high accuracy and robustness [21]. CIDR further uses an implicit imputation approach to alleviate the impact of dropouts in scRNA-seq data in a principled manner and identifies putative cell types using hierarchical clustering [22]. Other recent works are also proposed to cluster high dimensional single-cell data with gene regulatory networks [23], or ranking on shared nearest neighbors (SNN) [13].

However, there are still limitations in single-cell clustering to be addressed. First, even the state-of-the-art methods have achieved encouraging performance, the clustering quality can still be significantly improved for challenging tasks as shown in the Results Section. Second, most methods are designed for one of the continuous data and discrete UMI counts. There is demand for methods that can be applied to single-cell data created with different scRNA-seq technologies, such as discrete counts created with UMI based techniques and continuous data normalized to transcripts per million mapped reads (TPM)—if reads are only generated from one end of the transcript, or fragments per kilobase per million mapped reads (FPKM)—if reads cover the entire transcript [24]. Third, a method that can be generally applied to data of various sizes or dimensions is desirable. Nevertheless, most proposed methods cannot scale up well with various data dimensionality and with different dataset size. Last but equally important, most public packages cannot be easily installed due to system environment or software version problems. In addition, some of them require other dependency packages, and thus further inconveniences the use of these packages. Therefore, there is a demand for an easy-to-install and easy-to-use package.

Motivated by the above understanding, we propose in this paper a model-based algorithm named VPAC (Variational Projection for Accurate Clustering) of single-cell transcriptomic data. VPAC is a novel extension of the framework of probabilistic principal components analysis (PPCA) [25], which has been shown to be effective in dimensionality reduction for scRNA-seq data [26]. VPAC projects single-cell samples to a latent space, where the samples are constrained to follow a Gaussian mixture distribution [27]. With a coordinate ascent variational inference (CAVI) algorithm, VPAC can implement parameter estimation efficiently and steadily. Using five scRNA-seq datasets, we show that our model is not only superior to existing methods in the clustering of single-cell transcriptomic data, but also able to be applied to single-cell data of both discrete counts and normalized continuous data. Through comprehensive experiments, we further show the robustness of our model for various data dimensionality, different dataset size and different data sparsity, and the ability of our model to detect genes with strong unique signatures of a specific cell type.

## Methods

### Data collection

We collected a dataset of three T cell types (CD4+/CD25+ regulatory T cells, CD4+/CD45RA+/CD25- naive T cells and CD8+/CD45RA+ naïve cytotoxic T cells) from 10X Genomics [28]. The dataset measures the expression of 32,738 genes in 32,695 cells, which were enriched from fresh peripheral blood mononuclear cells (PBMCs) and sequenced by Illumina NextSeq 500 instrument. Clustering on this dataset was claimed as a challenging task by Z.Sun et al. [18] and T.Duan et al. [19]. We used this dataset to demonstrate the ability of our method to scale up well with various data dimensionality and with different dataset size on datasets from UMI based techniques. We also downloaded three preprocessed datasets of different scales used by Para-DPMM [19], which is the state-of-the-art method for clustering discrete UMI counts, to fairly evaluate the performance of our model.

A dataset of 561 cells derived from seven cell lines (A549, H1437, HCT116, IMR90, K562, GM12878, and H1) was downloaded from the NCBI Gene Expression Omnibus via accession GSE81861 [29]. Sequenced using the 101-bp paired-end protocol on the Illumina HiSeq 2000 platform, the dataset measures the expression of 55,186 DNA regions, and the expression was quantified as FPKM values. We used this dataset to demonstrate the robustness of our model for FPKM-normalized data of various dimensions. From the Single Cell Portal, we downloaded a plate-based scRNA-seq dataset of 24,649 genes in 27,998 cells from 12 clusters. The dataset was sequenced using a modified Smart-seq2 protocol, and the expression levels of genes were quantified as TPM values [30]. With this dataset, we further demonstrated the robustness of our model for TPM-normalized datasets of different size.

In order to illustrate the superior ability of our model to account for the sparsity of scRNA-seq data, we collected a dataset of 301 cells captured from 11 cell types using microfluidics. This dataset is publicly available from the NCBI Sequence Read Archive under accession SRP041736, and the quantification of gene expression levels in TPM for all 23730 genes in all samples was performed [31]. We also collected a Smart-seq2 sequenced dataset of 742 dendritic cells from six clusters with two batches from the Single Cell Portal to illustrate the application of our model [32, 33].

### The statistical model of VPAC

Let *N* be the number of single-cell samples, *D* the number of genes, and *M* the desired number of clusters. We assume that the gene expression vector x_**n**_ of cell *n* is generated from a projection of a latent *L*-dimensional vector **z**_**n**_ (*L* ≪ *D*). We use *n* = 1, …, *N* to index over samples, *i* = 1, …, *L* to index over latent dimensions, and *j* = 1, …, *M* to index over clusters in all derivations below. The distribution of x_n_ could be a complex high-dimensional distribution. We assume x_**n**_ follows a multivariate Gaussian distribution given **z**_**n**_, while the latent vector **z**_**n**_ follows a Gaussian mixture distribution. An intuitive understanding of this assumption is that, when projecting, constraining samples to follow a Gaussian mixture distribution in the latent space contributes to the accurate clustering. As a generative model, VPAC independently draw each sample x_**i**_ through the following process

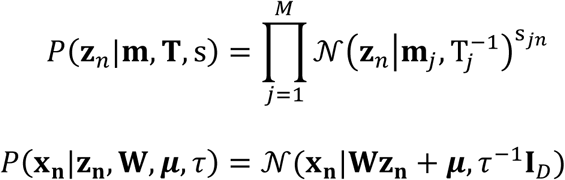

The binary latent variable s_***jn***_, which is given discrete distributions governed by **ρ**, describes which component in the mixture gives rise to the latent vector **z**_**n**_, i.e., if **z**_**n**_ is generated from component *j* then s_***jn***_ = 1, and s_***jn***_= 0 otherwise. We choose a Dirichlet distribution for the prior of **ρ**

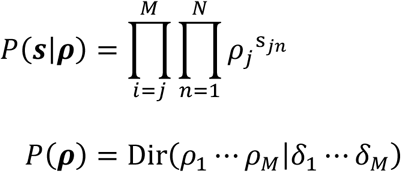

We complete the specification of Gaussian mixture distribution by introducing conjugate priors Gaussian distribution and Wishart distribution over the means **m** and precisions **T**. γ is a small and fixed parameter chosen to give a broad prior to **m**, while the degrees of freedom ***υ*** and scale matrix V give a broad prior to **T**

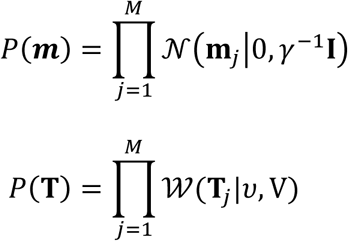

By introducing a hierarchical prior ***P***(**W**|α) over the projection matrix **W** in *P*(x_**n**_|**z**_**n**_, **W**, μ, τ), VPAC can automatically determine the appropriate dimensionality for the latent space to avoid discrete model selection. Each item of the *L*-dimensional vector **α** controls the corresponding column of the matrix **W** by playing a role as the precision of a Gaussian distribution

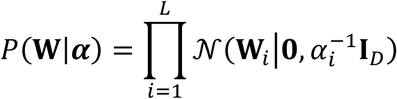

We again introduce broad priors over the parameters ***α, μ*** and *τ* to complete the specification of VPAC

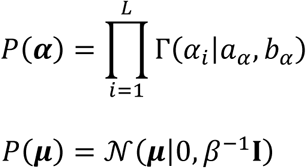

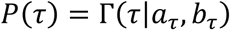

The graphical model representation of VPAC is shown in Figure 1. The broad priors introduced above are obtained by setting *a*_α_ = *b*_α_ = *a*_τ_ = *b*_τ_ = β = γ = 10^-3^. The initial parameters δ of the Dirichlet distribution are set as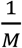. The joint distribution of all of the variables is given by

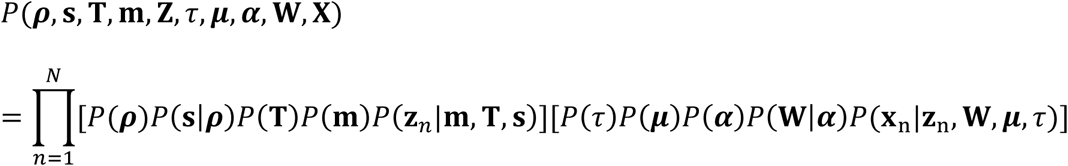

**Figure 1.**
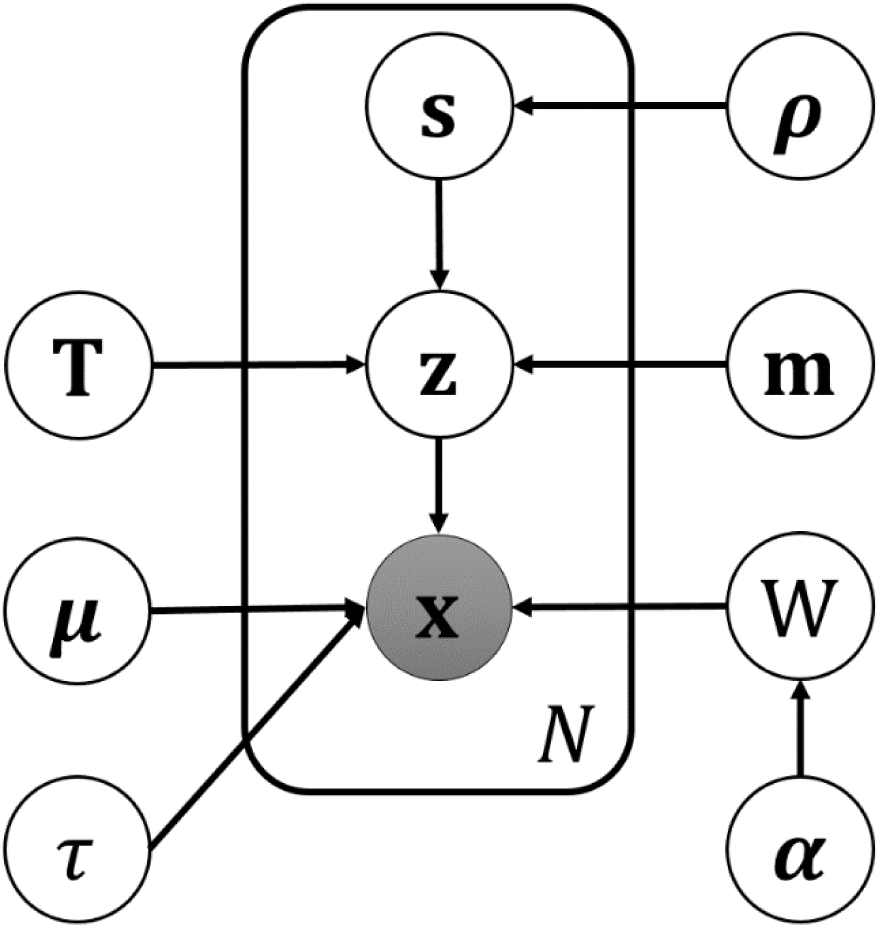
Representation of VPAC as a probabilistic graphical model. The observed variable **x** is shown by the shaded node, while the plate notation comprises a dataset of *N* independent observations together with the corresponding latent variables.

### Variational inference of parameters

We use variational methods to find a lower bound on *P*(**x**) because it is analytically intractable to directly evaluate *P*(**x**). With ***θ*** denoting the set of all parameters and latent variables in VPAC, we introduce an approximating distribution *Q*(***θ***) of the true posterior distribution. The log marginal likelihood is then given by

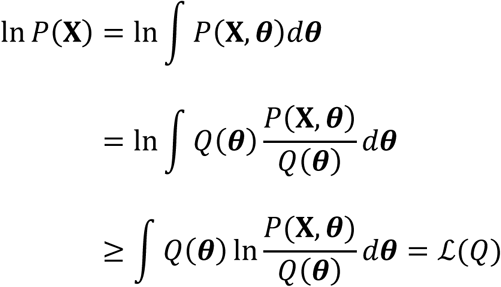

The function ℒ(*Q*) is the evidence lower bound (ELBO) on the true log marginal likelihood. The goal is to find a suitable *Q*(***θ***) to maximize the ELBO or minimize the Kullback-Leibler divergence between ℒ(*Q*) and the true log marginal likelihood

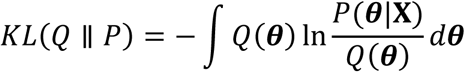

We assume the variational approximation is mean-field, i.e., *Q*(***θ***) = Π_*t*_ *Q*_*t*_(θ_*t*_). The corresponding factors are

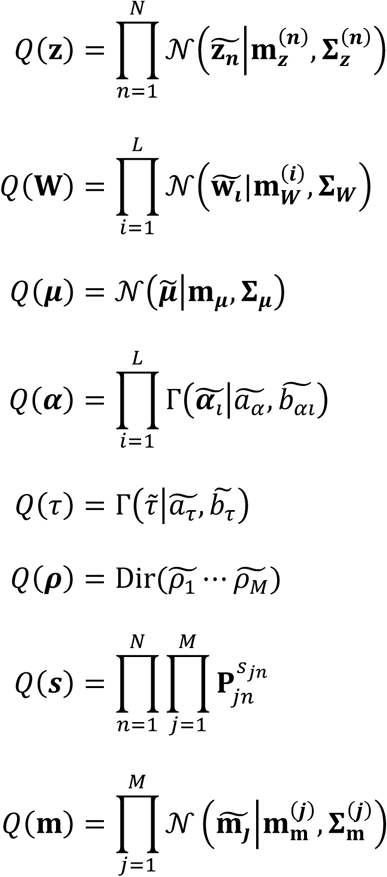

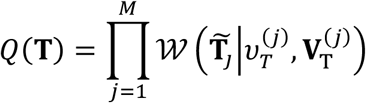

Because the proposed model is conjugate, we can derive a CAVI algorithm to update the variational parameters

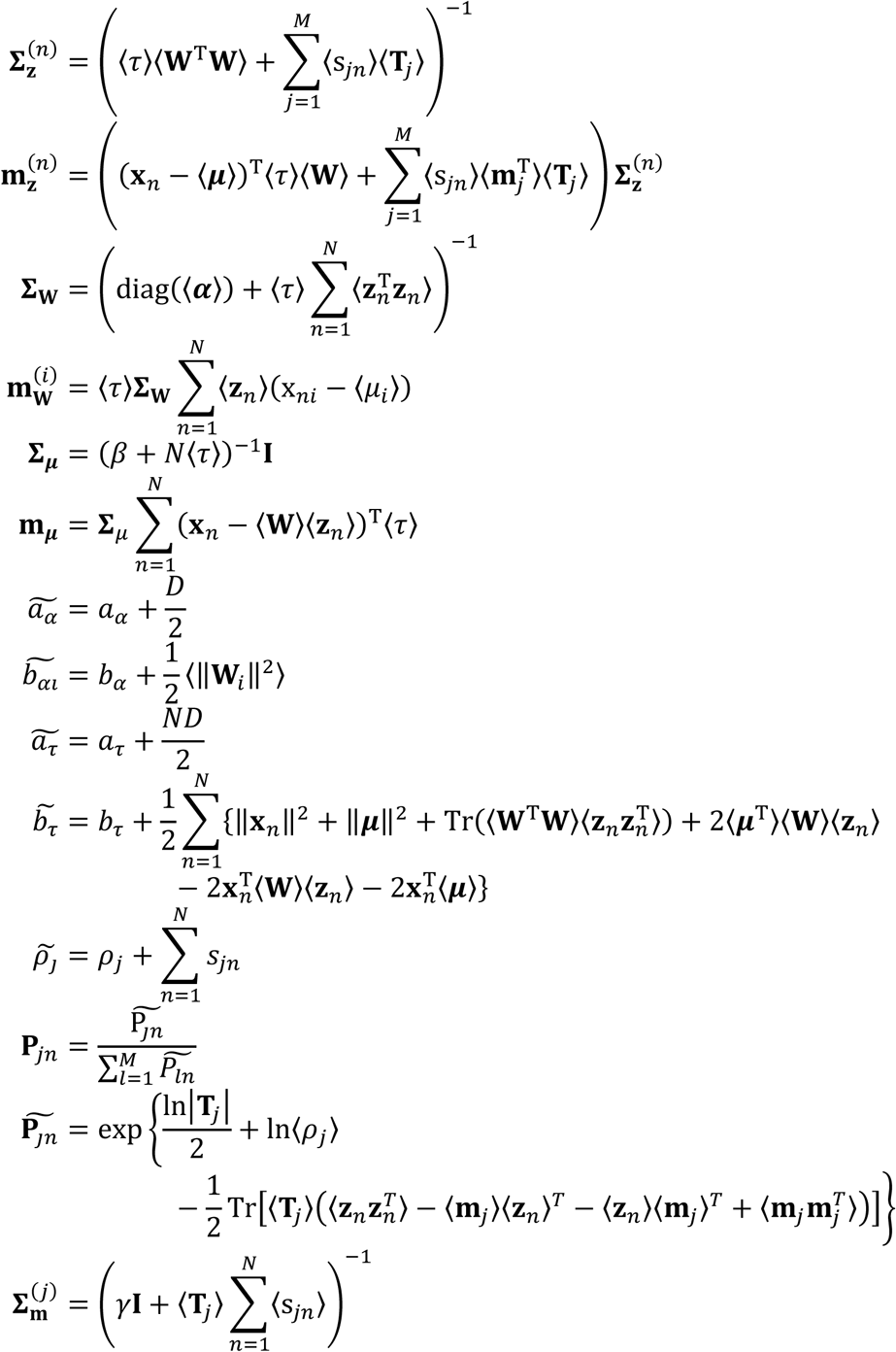

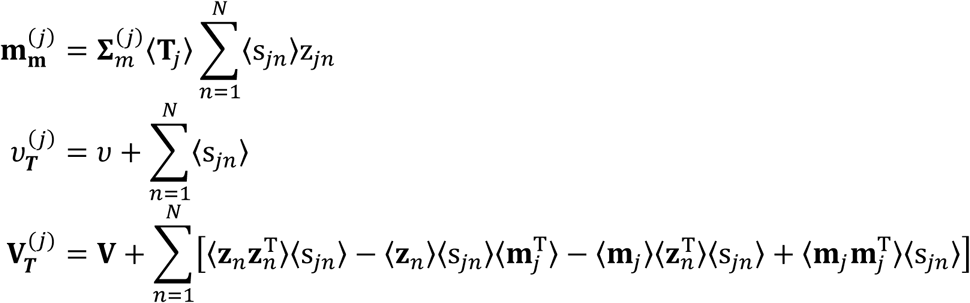

We implement VPAC in Python using common packages for data analysis (including Numpy, Scikit-learn and Scipy), without other dependency packages. Users can directly import our VPAC class and conduct clustering.

### Assessment of performance

We used adjusted rand index (ARI) [34] and normalized mutual information (NMI) [35] to assess the clustering performance, namely, the similarity between the clustering results and known cell types provided by their original references. Suppose T is the known cell types, C the predicted clustering results, *N* the total number of single-cell samples, *x*_*i*_ the number of samples clustered to the *i*-th cluster of C, *y*_***j***_ the number of samples belong to the *j*-th cell type of T, and ***n***_*i****j***_ the number of overlapping samples between the *i*-th cluster and the *j*-th cell type. ARI is computed as

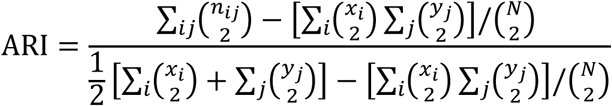

where ( ) denotes a binomial coefficient. By denoting the entropy of C and T as H(C) and H(T), respectively, and the mutual information between them as MI(C,T), NMI can be computed as

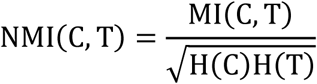

## Results

### VPAC accurately clusters scRNA-seq data

To verify the clustering performance of VPAC, we conducted each experiment five times and computed two widely used metrics, ARI and NMI. We compared the performance of our method with two baseline methods (using their default parameters), including Para_DPMM, the state-of-the-art method for clustering discrete UMI counts [19], and pcaReduce, another method combining dimension reduction with clustering [20].

In order to fairly evaluate the performance of our model, we compared the performance of VPAC with other two baseline methods on the three T cell datasets of different scales provided by T.Duan et al [19]. The small scale dataset (S-Set) consists of 1200 cells with the 1000 top variable genes, the medium scale dataset (M-Set) consists of 3000 cells with the 3000 top variable genes, and the large scale dataset (L-Set) consists of 6000 cells with the 5000 top variable genes. As shown in Table 1, our method consistently outperforms the two baseline methods for a large margin on all the three datasets, where it achieves average 11.3% improvement on ARI and 8.1% improvement on NMI compared to Para_DPMM, even the clustering on these datasets was claimed as a challenging task [18, 19]. In addition, our method consistently achieves stable performance, which demonstrates its higher robustness than the baseline methods. In terms of training efficiency, taking the L-Set as an example, VPAC cost 241 seconds, Para_DPMM cost 1000 seconds and pcaReduce cost 37 seconds on average. It is worth noting that the default minimal training time of Para_DPMM is 1000 seconds, which slightly improves the performance of Para_DPMM shown in the original research.

**Table 1.** Performance comparison on three datasets of different scales.

| Dataset | S-Set |  | M-Set |  | L-Set |  |
| --- | --- | --- | --- | --- | --- | --- |
| Model | ARI | NMI | ARI | NMI | ARI | NMI |
| VPAC | **0.769 ±0.000** | **0.759 ±0.000** | **0.765 ±0.000** | **0.765 ±0.000** | **0.779 ±0.000** | **0.769 ±0.000** |
| Para_DPMM | 0.675 ±0.004 | 0.696 ±0.007 | 0.704 ±0.000 | 0.713 ±0.001 | 0.700 ±0.002 | 0.711 ±0.002 |
| pcaReduce | 0.290 ±0.005 | 0.289 ±0.004 | 0.279 ±0.007 | 0.252 ±0.007 | 0.311 ±0.023 | 0.309 ±0.044 |

### VPAC scales up well with various data dimensionality

To illustrate the robustness of our model for various data dimensionality, we selected different numbers of top variable genes (features) based on their standard deviations across the cell transcriptome profiles. We first used the dataset of 32,695 cells enriched from PBMCs, which is the full version of the three preprocessed datasets, to demonstrate the superior performance of VPAC for discrete UMI counts of various dimensionality. As illustrated in Figure 2a, our method significantly outperforms the two baseline methods and achieves consistent performance across datasets of different numbers of features. Note that pcaReduce raised an error when training the dataset with 30k features even the memory is sufficient. We further evaluated the robustness of our model for FPKM-normalized data using a dataset of 561 cells derived from seven cell lines. As shown in Figure 2b, different with the performance for discrete UMI counts, the performance of pcaReduce is superior to that of Para_DPMM, and the performance of these two baseline methods are obviously unstable, while VPAC, again, outperforms the baseline methods and achieves consistent performance across datasets of different numbers of features. The results demonstrate that VPAC can scale up well with various data dimensionality whether the scRNA-seq data is discrete or continuous.

**Figure 2.**
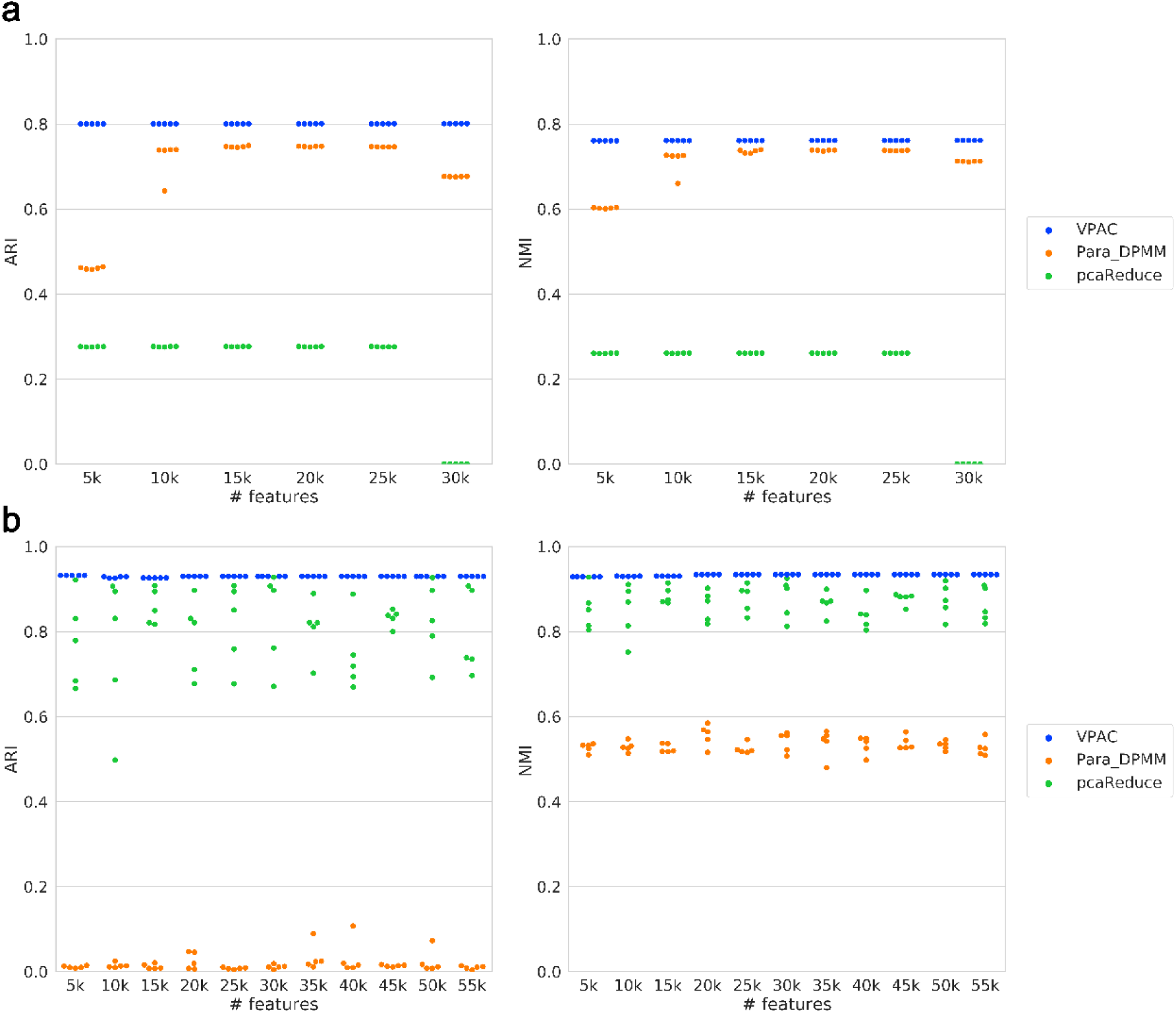
Performance comparison on datasets of various dimensionality. (a) The performance on discrete UMI counts. (b) The performance on continuous FPKM-normalized data.

### VPAC scales up well with different dataset size

Different scRNA-seq technologies can be used to quantify transcriptomes for different numbers of cells. In order to demonstrate the robustness of our model for different dataset size, we randomly selected different proportions of single-cell samples from the whole cell population. Besides, we selected top 90% variable genes (features) to avoid errors raised by pcaReduce. We again used the cells enriched from PBMCs to illustrate the superior performance of VPAC for discrete UMI counts of different dataset size. As illustrated in Figure 3a, our method significantly outperforms the two baseline methods and achieves superior performance across datasets of different numbers of samples, while the performance of Para_DPMM deteriorates as the number of samples increased. We further illustrated the robustness of our model for TPM-normalized data using a dataset of 24649 cells from 12 clusters. As shown in Figure 3b, VPAC is still superior to the baseline methods and achieves more stable performance across datasets of different numbers of samples, while the performance of pcaReduce is still unstable even outperforms Para_DPMM again. The results above demonstrate that VPAC can also scale up well with different dataset size whether the scRNA-seq data is discrete or continuous.

**Figure 3.**
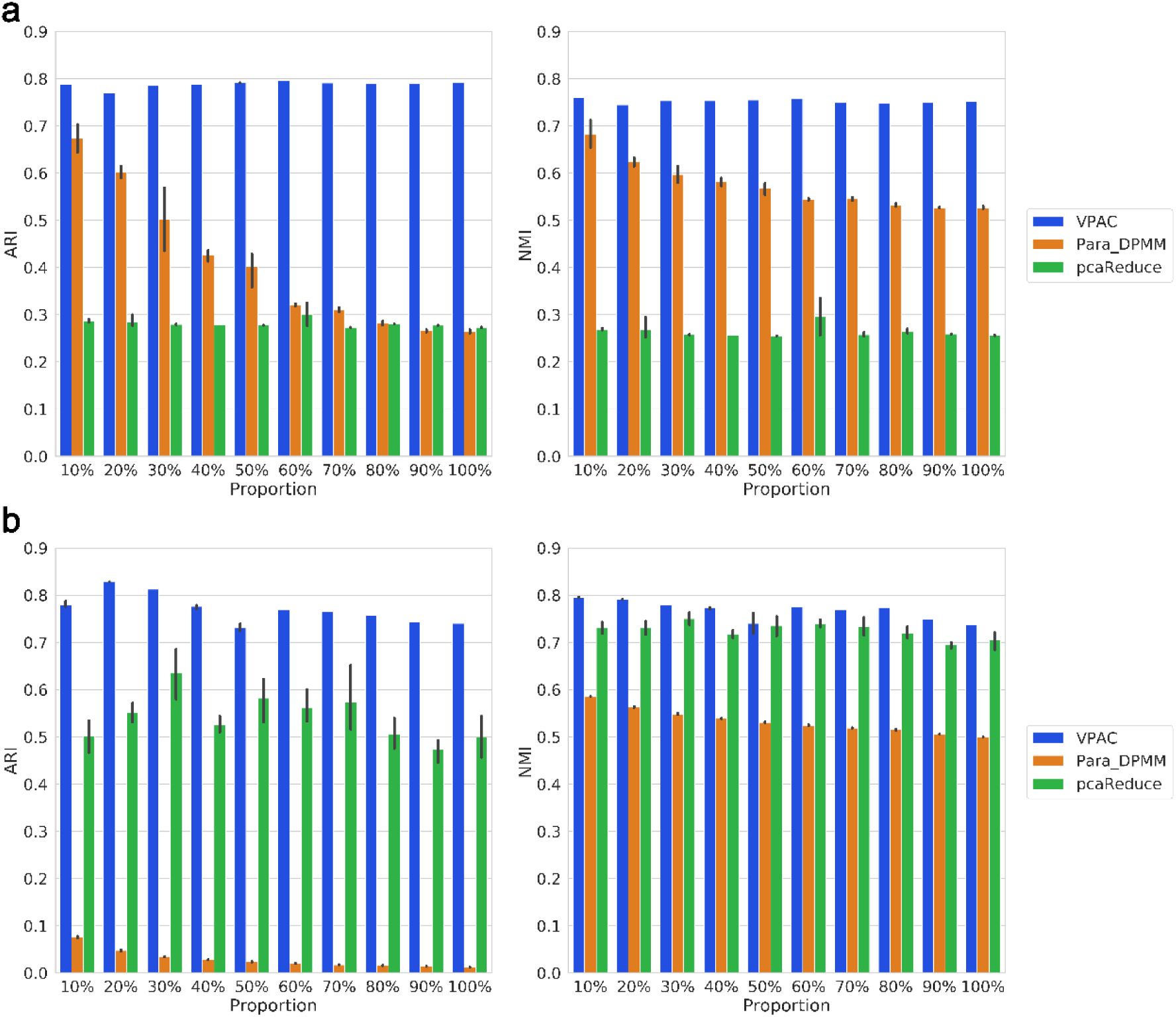
Performance comparison on datasets of different size. (a) The performance on discrete UMI counts. (b) The performance on continuous TPM-normalized data.

### VPAC scales up well with different data sparsity

Single-cell gene expression data suffer from the high sparsity problem because they contain an abundance of dropout events that lead to zero expression measurements. In order to illustrate the superior ability of our model to account for the sparsity of scRNA-seq data, we randomly zeroed different proportions of non-zero items five times on a TPM-normalized dataset of 301 cells with about 2.7 × 10^5^ reads per cell [31]. As illustrated in Figure 4, VPAC is superior to the baseline methods and achieves more stable performance. The performance of both VPAC and pcaReduce obviously deteriorate when we set 80% of the non-zero expression data to zero, while Para_DPMM again fails on clustering the TPM-normalized dataset. The results demonstrate that our model has the ability to account for the sparsity of scRNA-seq data, and thus benefits the clustering tasks with the exponential growth of the Drop-seq based single-cell transcriptomic data.

**Figure 4.**
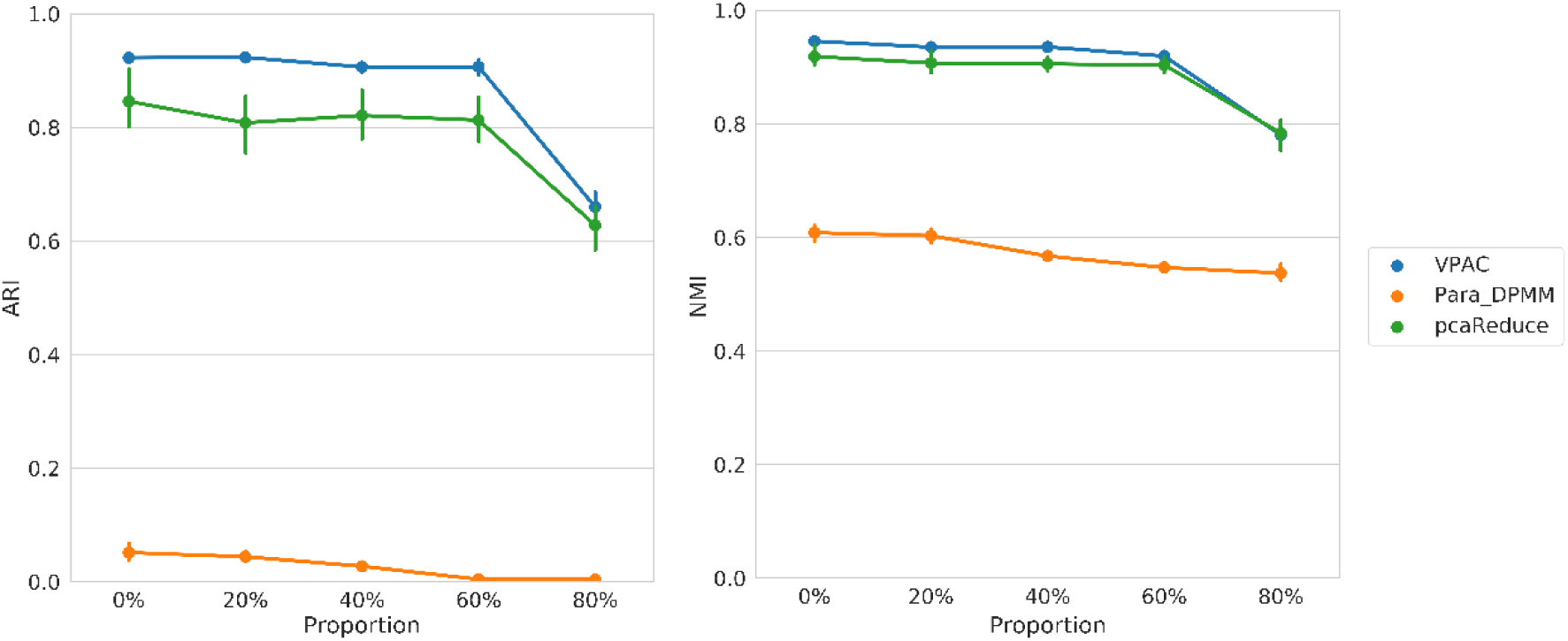
Performance comparison on datasets of different sparsity.

### Applications of VPAC

To demonstrate potential applications of VPAC, we collected a dataset of 742 dendritic cells from six clusters [32, 33]. By setting the number of clusters to 6 and the dimensionality of latent space to 10, VPAC achieved an ARI of 0.876 and an NMI of 0.898. Actually, the items of α with large values in VPAC will result in ‘switching off’ the corresponding column of the matrix **W**, and thus provide guidance for determining the appropriate dimensionality for the latent space. As shown in Figure 5, the visualization of the projection matrix **W** (the left one) indicates there are four principal directions in the latent space, while the visualization of the projection matrix obtained by classical PCA (the right one) cannot effectively reveal the number of principal directions. Therefore, by setting the dimensionality of latent space to 5 (in practice we have found that one more dimensionality, which may contain some additional information, tends to give better results), VPAC achieved an ARI of 0.875 and an NMI of 0.903, which are approximately equal to the results with the dimensionality of latent space setting to 10, which means the only one parameter we should determine for VPAC is the number of clusters.

**Figure 5.**
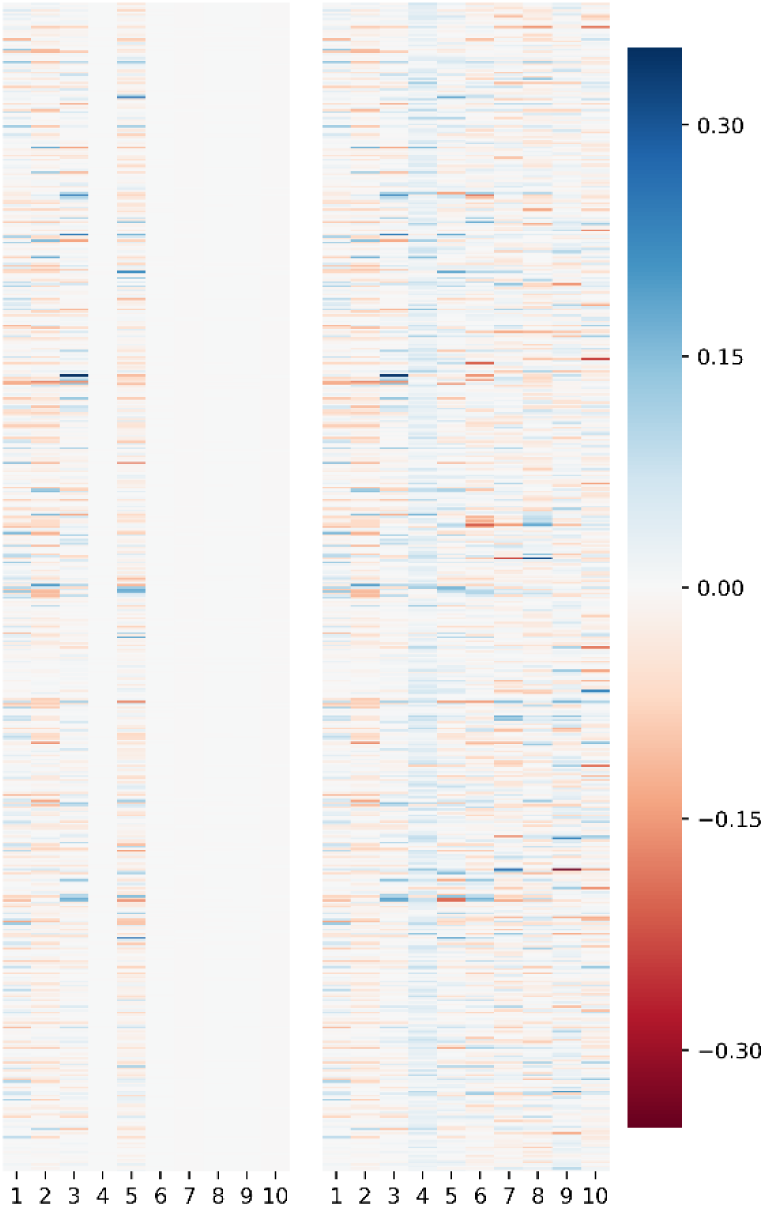
Visualization of the projection matrix W of VPAC (the left one) and that of classical PCA (the right one).

We used the t-SNE algorithm to visualize the dimension-reduced data in latent space of VPAC by projecting the data into a two-dimensional space so that certain hidden structures can be depicted intuitively. Note that t-SNE is a visualization tool, and it is not intended to be used for clustering scRNA-seq data [18]. As shown in Figure 6a, VPAC accurately clustered different cell types accounting for cells from different batches. It is worth noting that VPAC intended to cluster DC2 and DC3 cell types together while inferring a small cluster (in the bottom) containing some individual cells. According to the original research for this dataset, it is reasonable to cluster DC2 and DC3 together because both of them correspond to new subdivisions of the CD1C/BDCA-1^+^cDC2, while DC1 corresponds to the cross-presenting CD141/BDCA- 3^+^ cDC1, DC4 corresponds to CD1C ^−^CD141^−^CD11C^+^ DC, DC5 is a unique DC subtype, AS DCs, and DC6 corresponds to the interferon-producing pDC [32]. With this understanding, we set the number of clusters to 5. VPAC successfully clustered the DC2/3 cluster and achieved an ARI of 0.892 and an NMI of 0.943. In addition, as illustrated in Figure 6(b), VPAC accounted for batch effects even there are still very few samples wrongly clustered, which means that our method is robust to scRNA-seq data from different batches.

**Figure 6.**
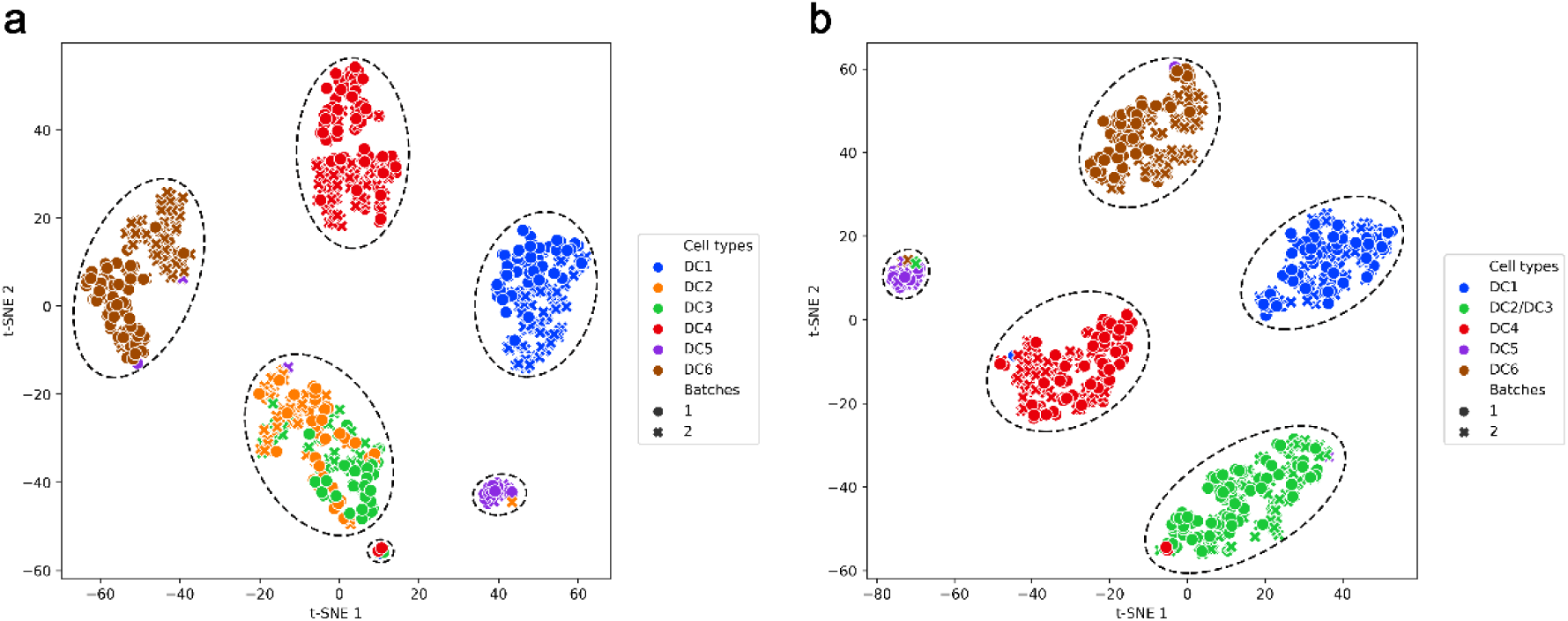
Visualization of the dendritic cells in latent space inferred by VPAC using t-SNE. The dendritic cells are colored by cell-type labels provided by the original study, different shapes of points represent different experimental batches, and the dashed circles represent potential clusters inferred by VPAC with setting of the number of clusters to **(a)** 6, and **(b)** 5.

We further conducted gene co-expression network analysis to identify which genes have a tendency to show a coordinated expression pattern of the DC2/3 cluster inferred by VPAC. By calculating the Pearson correlation coefficient (PCC) of each gene pairs, we plotted a co-expression network with a threshold 0.4 of PCC [36]. As illustrated in Figure 7, the three genes, namely S100A9, S100A8 and CD14, with the highest degree of connectivity are expected to be drivers required for signaling pathways of essential functions. It is worth noting that, in the original research, these three genes were claimed as acute and chronic inflammatory genes that play as a strong unique signature to distinguish the DC2/3 cluster, which means our method has the ability to reveal the genes with strong unique signatures of a specific cell type.

**Figure 7.**
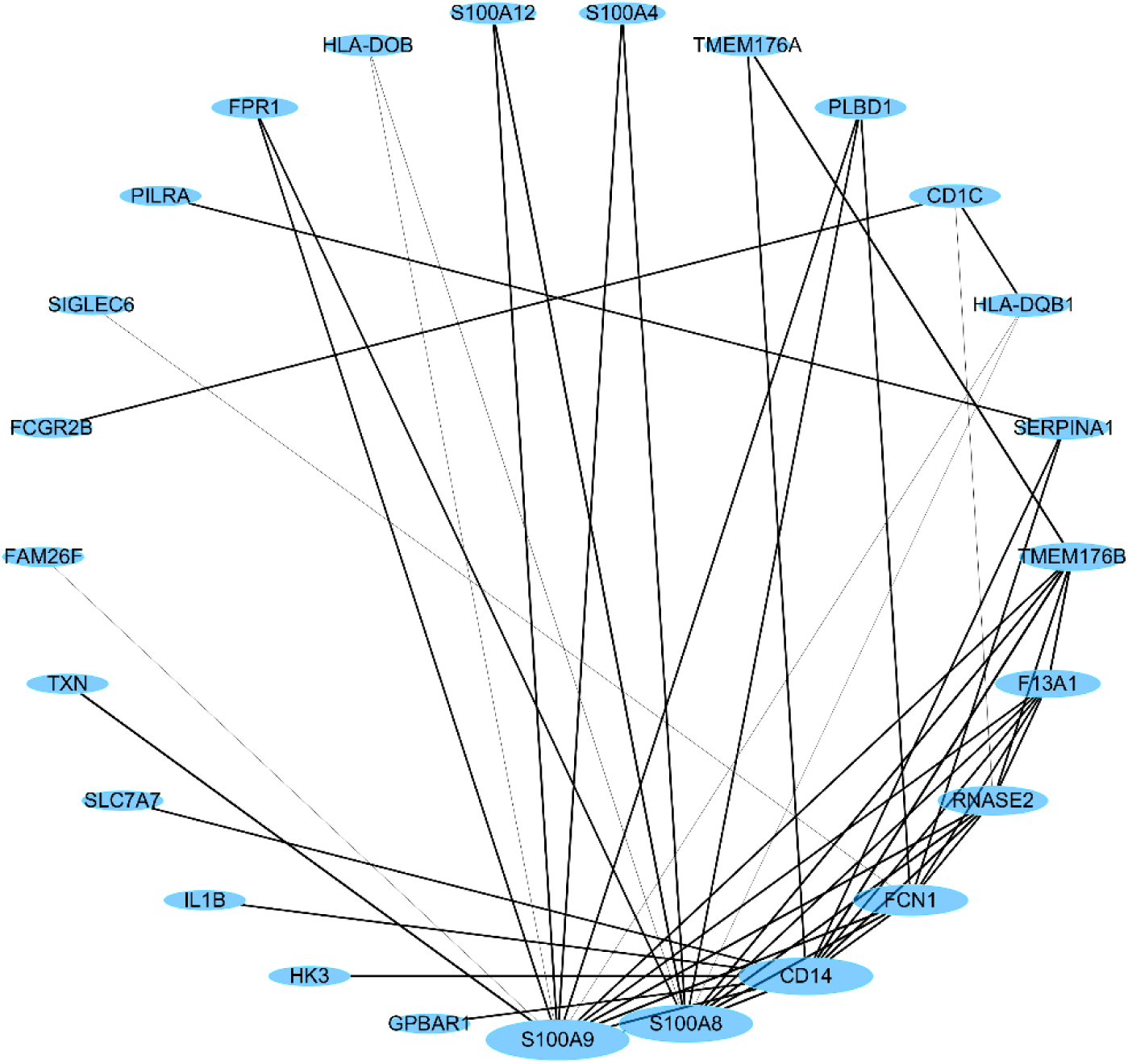
The co-expression network of the DC2/3 cluster inferred by VPAC.

It is worth noting that VPAC successfully clustered the novel DC5 samples detected by the original research. In order to study the potential mechanism implied in VPAC, we further analyzed the samples in the latent space inferred by VPAC. As shown in Figure 8, the fifth items in latent vectors of the DC5 cluster are much larger than that of other clusters, which means that the fifth direction of the latent space can effectively distinguish the DC5 cluster. We sorted the absolute values of the fifth column in projection matrix **W**, which are the absolute weights of linear combinations of genes, and found that the two genes with the highest absolute weights (0.290 and 0.288) are AXL and SIGLEC6, which have much higher absolute weight than the successive gene CX3CR1 with a weight of 0.225. Interestingly, AXL and SIGLEC6 serve as unique markers for the population emerged from the unbiased cluster analysis (cluster DC5) according to the original literature, which further demonstrates that our method has the inherent ability to detect genes with strong unique signatures of a specific cell type while accurately clustering.

**Figure 8.**
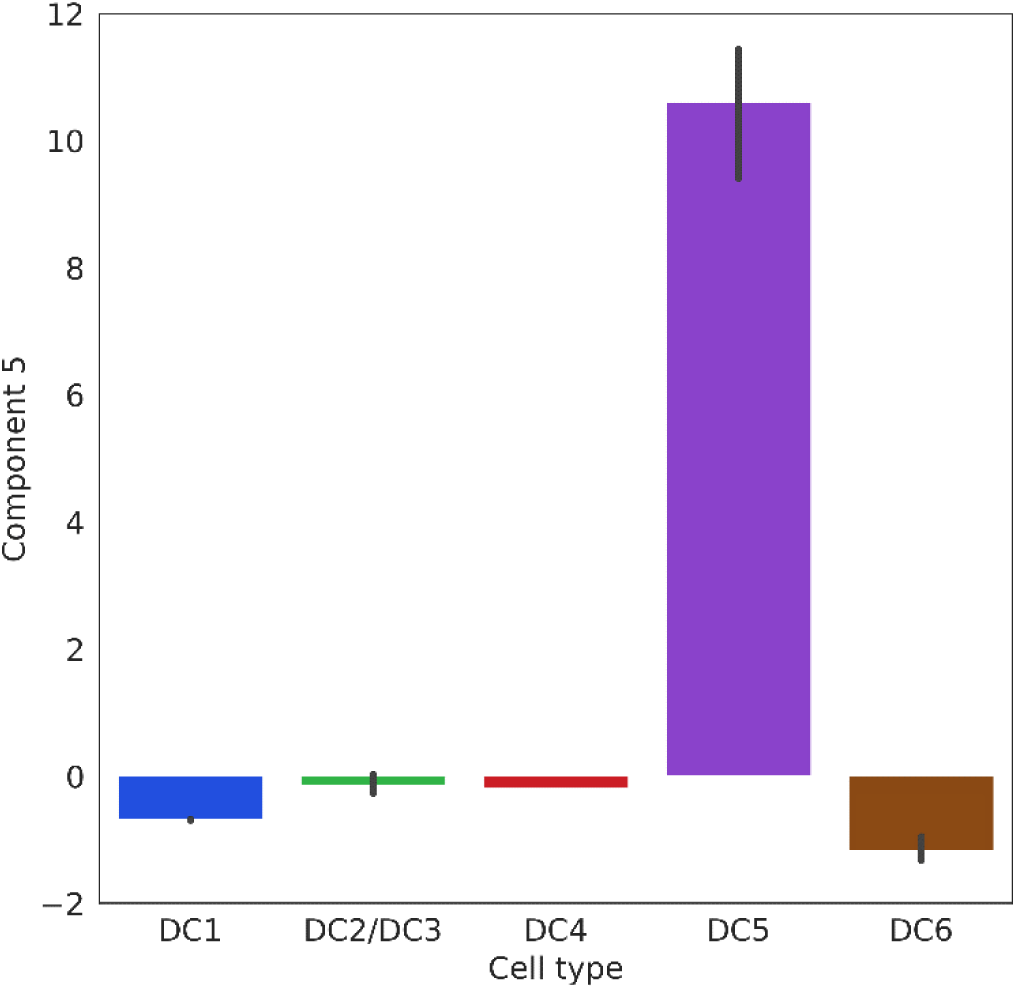
The values of fifth items in the latent vectors inferred by VPAC.

## Discussion

scRNA-Seq technologies have advanced rapidly in recent years and enable the quantitative characterization of cell types based on transcriptome profiles. VPAC is proposed to identify putative cell types using unsupervised clustering. The superiority of our method over other baseline methods, such as Para_DPMM and pcaReduce, is mainly attributed to the variational projection while constraining single-cell samples to follow a Gaussian mixture distribution in the latent space. In addition, with the hierarchical prior over the projection matrix, VPAC can automatically determine the appropriate dimensionality for the latent space to avoid discrete model selection, which means the only one parameter we should determine for VPAC is the number of clusters. The comprehensive experiments demonstrate the generality and robustness of our model.

Our model can certainly be improved in some aspects. First, a more complex non-linear projection can be introduced to better model the scRNA-seq data. Second, more constraints or assumptions can be included in our model considering the characteristics of scRNA-seq data, such as the high sparsity. Recent studies have shown the introduction of zero-inflated assumption can effectively model the dropout evens of single-cell data [26, 37], which may also improve the performance of our model. Third, our model can be extended to incorporate other types of functional genomics data such as chromatin accessibility. For example, in the literature, the method of coupled nonnegative matrix factorizations performs clustering by the integrative analysis of scRNA-seq and single-cell ATAC-sequencing data [38]. Finally, the performance and efficiency of the model may be further improved by parameters inference using stochastic optimization and deep neural networks, which have been shown to be effective in statistical models for single-cell data analysis [39, 40].

## Conclusions

We have proposed a model-based algorithm, named VPAC, for accurate clustering of single-cell transcriptomic data through variational projection. Benefitting from the variational projection while constraining single-cell samples to follow a Gaussian mixture distribution in the latent space, VPAC is superior to existing methods in the clustering of datasets of discrete counts, normalized continuous data, various data dimensionality, different dataset size and different data sparsity. We have further demonstrated the ability of VPAC to detect genes with strong unique signatures of a specific cell type, which may shed light on the studies in system biology. Eventually, with the explosive growth of scRNA-seq data, we expect that such a statistical approach will provide us superior performance and be widely applicable.

## Declarations

### Abbreviations

scRNA-seq: : single-cell RNA-sequencing
UMI: : unique molecular identifiers
t-SNE: : t-Distributed Stochastic Neighbor Embedding
PCA: : principal component analysis
SNN: : shared nearest neighbors
TPM: : transcripts per million mapped reads
FPKM: : fragments per kilobase per million mapped reads
PPCA: : probabilistic principal components analysis
CAVI: : coordinate ascent variational inference
PBMCs: : peripheral blood mononuclear cells
ELBO: : evidence lower bound
ARI: : adjusted rand index
NMI: : normalized mutual information
PCC: : Pearson correlation coefficient

### Ethics approval and consent to participate

Not applicable.

### Consent for publication

Not applicable.

### Availability of data and material

The datasets supporting the conclusions of this article are publicly available from 10xgenomics (https://support.10xgenomics.com/single-cell-gene-expression/datasets), the NCBI Gene Expression Omnibus (https://www.ncbi.nlm.nih.gov/geo/), the NCBI Sequence Read Archive (http://www.ncbi.nlm.nih.gov/Traces/sra/), and the Single Cell Portal (https://portals.broadinstitute.org/single_cell).

### Competing interests

The authors declare that they have no competing interests.

### Funding

This research was partially supported by the National Key Research and Development Program of China (No. 2018YFC0910404), the National Natural Science Foundation of China (Nos. 61873141, 61721003, 61573207, 71871019 and 71471016), and the Tsinghua-Fuzhou Institute for Data Technology.

### Authors’ contributions

RJ designed the research. SC and KH designed and implemented the models. SC and HC collected data and analyzed the results. SC, KH, HC and RJ wrote the manuscript. All authors read and confirmed the manuscript.

## Acknowledgements

We thank Yong Wang for his helpful discussions. Rui Jiang is a RONG professor at the Institute for Data Science, Tsinghua University.

## References

1. Tang F, Barbacioru C, Wang Y, Nordman E, Lee C, Xu N, Wang X, Bodeau J, Tuch BB, Siddiqui A et al. mRNA-Seq whole-transcriptome analysis of a single cell. Nat Methods. 2009; 6(5):377–382.

2. Hedlund E, Deng Q. Single-cell RNA sequencing: Technical advancements and biological applications. Mol Aspects Med. 2018; 59:36–46.

3. Picelli S, Bjorklund AK, Faridani OR, Sagasser S, Winberg G, Sandberg R. Smart-seq2 for sensitive full-length transcriptome profiling in single cells. Nat Methods. 2013; 10(11):1096–1098.

4. Jaitin DA, Kenigsberg E, Keren-Shaul H, Elefant N, Paul F, Zaretsky I, Mildner A, Cohen N, Jung S, Tanay A et al. Massively parallel single-cell RNA-seq for marker-free decomposition of tissues into cell types. Science. 2014; 343(6172):776–779.

5. Macosko EZ, Basu A, Satija R, Nemesh J, Shekhar K, Goldman M, Tirosh I, Bialas AR, Kamitaki N, Martersteck EM et al. Highly Parallel Genome-wide Expression Profiling of Individual Cells Using Nanoliter Droplets. Cell. 2015; 161(5):1202–1214.

6. Ziegenhain C, Vieth B, Parekh S, Reinius B, Guillaumet-Adkins A, Smets M, Leonhardt H, Heyn H, Hellmann I, Enard W. Comparative Analysis of Single-Cell RNA Sequencing Methods. Mol Cell. 2017; 65(4):631–643 e634.

7. Segerstolpe A, Palasantza A, Eliasson P, Andersson EM, Andreasson AC, Sun X, Picelli S, Sabirsh A, Clausen M, Bjursell MK et al. Single-Cell Transcriptome Profiling of Human Pancreatic Islets in Health and Type 2 Diabetes. Cell Metab. 2016; 24(4):593–607.

8. Grun D, Lyubimova A, Kester L, Wiebrands K, Basak O, Sasaki N, Clevers H, van Oudenaarden A. Single-cell messenger RNA sequencing reveals rare intestinal cell types. Nature. 2015; 525(7568):251–255.

9. Zeisel A, Munoz-Manchado AB, Codeluppi S, Lonnerberg P, La Manno G, Jureus A, Marques S, Munguba H, He L, Betsholtz C et al. Brain structure. Cell types in the mouse cortex and hippocampus revealed by single-cell RNAseq. Science. 2015; 347(6226):1138–1142.

10. Scialdone A, Natarajan KN, Saraiva LR, Proserpio V, Teichmann SA, Stegle O, Marioni JC, Buettner F. Computational assignment of cell-cycle stage from single-cell transcriptome data. Methods. 2015; 85:54–61.

11. Jaitin DA, Weiner A, Yofe I, Lara-Astiaso D, Keren-Shaul H, David E, Salame TM, Tanay A, van Oudenaarden A, Amit I. Dissecting Immune Circuits by Linking CRISPR-Pooled Screens with Single-Cell RNA-Seq. Cell. 2016; 167(7):1883–1896 e1815.

12. Xue Z, Huang K, Cai C, Cai L, Jiang CY, Feng Y, Liu Z, Zeng Q, Cheng L, Sun YE et al. Genetic programs in human and mouse early embryos revealed by single-cell RNA sequencing. Nature. 2013; 500(7464):593–597.

13. Satija R, Farrell JA, Gennert D, Schier AF, Regev A. Spatial reconstruction of single-cell gene expression data. Nat Biotechnol. 2015; 33(5):495–502.

14. Marco E, Karp RL, Guo G, Robson P, Hart AH, Trippa L, Yuan GC. Bifurcation analysis of single-cell gene expression data reveals epigenetic landscape. Proc Natl Acad Sci U S A. 2014; 111(52):E5643–5650.

15. Treutlein B, Brownfield DG, Wu AR, Neff NF, Mantalas GL, Espinoza FH, Desai TJ, Krasnow MA, Quake SR. Reconstructing lineage hierarchies of the distal lung epithelium using single-cell RNA-seq. Nature. 2014; 509(7500):371–375.

16. Stegle O, Teichmann SA, Marioni JC. Computational and analytical challenges in single-cell transcriptomics. Nat Rev Genet. 2015; 16(3):133–145.

17. duVerle DA, Yotsukura S, Nomura S, Aburatani H, Tsuda K. CellTree: an R/bioconductor package to infer the hierarchical structure of cell populations from single-cell RNA-seq data. BMC Bioinformatics. 2016; 17(1):363.

18. Sun Z, Wang T, Deng K, Wang XF, Lafyatis R, Ding Y, Hu M, Chen W. DIMM-SC: a Dirichlet mixture model for clustering droplet-based single cell transcriptomic data. Bioinformatics. 2018; 34(1):139–146.

19. Duan T, Pinto JP, Xie X. Parallel Clustering of Single Cell Transcriptomic Data with Split-Merge Sampling on Dirichlet Process Mixtures. Bioinformatics. 2018.

20. Zurauskiene J, Yau C. pcaReduce: hierarchical clustering of single cell transcriptional profiles. BMC Bioinformatics. 2016; 17:140.

21. Kiselev VY, Kirschner K, Schaub MT, Andrews T, Yiu A, Chandra T, Natarajan KN, Reik W, Barahona M, Green AR et al. SC3: consensus clustering of single-cell RNA-seq data. Nat Methods. 2017; 14(5):483–486.

22. Lin P, Troup M, Ho JW. CIDR: Ultrafast and accurate clustering through imputation for single-cell RNA-seq data. Genome Biol. 2017; 18(1):59.

23. Aibar S, Gonzalez-Blas CB, Moerman T, Huynh-Thu VA, Imrichova H, Hulselmans G, Rambow F, Marine JC, Geurts P, Aerts J et al. SCENIC: single-cell regulatory network inference and clustering. Nat Methods. 2017; 14(11):1083–1086.

24. Grun D, van Oudenaarden A. Design and Analysis of Single-Cell Sequencing Experiments. Cell. 2015; 163(4):799–810.

25. Tipping ME, Bishop CMJJotRSSSB. Probabilistic principal component analysis. Journal of the Royal Statistical Society. 1999; 61(3):611–622.

26. Pierson E, Yau C. ZIFA: Dimensionality reduction for zero-inflated single-cell gene expression analysis. Genome Biol. 2015; 16:241.

27. Corduneanu A, Bishop CM: Variational Bayesian model selection for mixture distributions. In: Artificial intelligence and Statistics: 2001. >Morgan Kaufmann Waltham, MA: 27–34.

28. Zheng GX, Terry JM, Belgrader P, Ryvkin P, Bent ZW, Wilson R, Ziraldo SB, Wheeler TD, McDermott GP, Zhu J et al. Massively parallel digital transcriptional profiling of single cells. Nat Commun. 2017; 8:14049.

29. Li H, Courtois ET, Sengupta D, Tan Y, Chen KH, Goh JJL, Kong SL, Chua C, Hon LK, Tan WS et al. Reference component analysis of single-cell transcriptomes celucidates cellular heterogeneity in human colorectal tumors. Nat Genet. 2017; 49(5):708–718.

30. Biton M, Haber AL, Rogel N, Burgin G, Beyaz S, Schnell A, Ashenberg O, Su CW, Smillie C, Shekhar K et al. T Helper Cell Cytokines Modulate Intestinal Stem Cell Renewal and Differentiation. Cell. 2018; 175(5):1307–1320 e1322.

31. Pollen AA, Nowakowski TJ, Shuga J, Wang X, Leyrat AA, Lui JH, Li N, Szpankowski L, Fowler B, Chen P et al. Low-coverage single-cell mRNA sequencing reveals cellular heterogeneity and activated signaling pathways in developing cerebral cortex. Nat Biotechnol. 2014; 32(10):1053–1058.

32. Villani AC, Satija R, Reynolds G, Sarkizova S, Shekhar K, Fletcher J, Griesbeck M, Butler A, Zheng S, Lazo S et al. Single-cell RNA-seq reveals new types of human blood dendritic cells, monocytes, and progenitors. Science. 2017; 356(6335).

33. Hua K, Zhang X. A case study on the detailed reproducibility of a human cell atlas project. 2018:467993.

34. Hubert L, Arabie PJJoc. Comparing partitions. Journal of classification. 1985; 2(1):193–218.

35. Strehl A, Ghosh JJJomlr. Cluster ensembles---a knowledge reuse framework for combining multiple partitions. 2002; 3(Dec):583–617.

36. Shannon P, Markiel A, Ozier O, Baliga NS, Wang JT, Ramage D, Amin N, Schwikowski B, Ideker T. Cytoscape: a software environment for integrated models of biomolecular interaction networks. Genome Res. 2003; 13(11):2498–2504.

37. Ferreira PF, Carvalho AM, Vinga S. Scalable probabilistic matrix factorization for single-cell RNA-seq analysis. bioRxiv. 2018:496810.

38. Duren Z, Chen X, Zamanighomi M, Zeng W, Satpathy AT, Chang HY, Wang Y, Wong WH. Integrative analysis of single-cell genomics data by coupled nonnegative matrix factorizations. Proc Natl Acad Sci U S A. 2018; 115(30):7723–7728.

39. Lopez R, Regier J, Cole MB, Jordan MI, Yosef N. Deep generative modeling for single-cell transcriptomics. Nat Methods. 2018; 15(12):1053–1058.

40. Ding J, Condon A, Shah SP. Interpretable dimensionality reduction of single cell transcriptome data with deep generative models. Nat Commun. 2018; 9(1):2002.

